# Biophysical characterization of the inactivation of *E. coli* transketolase by aqueous co-solvents

**DOI:** 10.1101/2020.06.09.140988

**Authors:** Phattaraporn Morris, Ribia García-Arrazola, Leonardo Rios-Solis, Paul A. Dalby

**Affiliations:** Department of Biochemical Engineering, University College London, Bernard Katz Building, Gower Street, London WC1E 6BT, UK; Chemical Metrology and Biometry Department, National Institute of Metrology, 3/4-5 Moo 3, Klong 5, Klong Luang, Pathumthani 12120, Thailand; Institute for Bioengineering, School of Engineering, University of Edinburgh, Edinburgh, EH9 3JL, UK; Centre for Synthetic and Systems Biology (SynthSys), University of Edinburgh, King’s Buildings, Edinburgh, United Kingdom, EH9 3JL, UK

**Keywords:** transketolase, co-solvents, inactivation, local unfolding, aggregation

## Abstract

Transketolase (TK) has been previously engineered, using semi-rational directed evolution and substrate walking, to accept increasingly aliphatic, cyclic and then aromatic substrates. This has ultimately led to the poor water solubility of new substrates, as a potential bottleneck to further exploitation of this enzyme in biocatalysis. Here we used a range of biophysical studies to characterise the response of both *E. coli* apo- and holo-TK activity and structure to a range of commonly used polar organic co-solvents: acetonitrile (MeCN), *n*- butanol (nBuOH), ethyl acetate (EToAc), isopropanol (iPrOH), and tetrahydrofuran (THF). The mechanism of enzyme deactivation was found to be predominantly via solvent-induced local unfolding. Holo-TK is thermodynamically more stable than apo-TK and yet for four of the five co-solvents it retained less activity than apo-TK after exposure to organic solvents, indicating that solvent tolerance was not correlated to global conformational stability. The co-solvent concentrations required for complete enzyme inactivation was inversely proportional to co-solvent log(P), while the unfolding rate was directly proportional, indicating that the solvents interact with and partially unfold the enzyme through hydrophobic contacts. Aggregation was not found to be the driving mechanism of enzyme inactivation, but was in some cases an additional impact of solvent-induced local or global unfolding.

TK was found to be tolerant to 15% (v/v) iPrOH, 10% (v/v) MeCN, or 6% (v/v) nBuOH over 3 hours. This work indicates that future attempts to engineer the enzyme to better tolerate co-solvents should focus on increasing the stability of the protein to local unfolding, particularly in and around the cofactor-binding loops.

## 1. Introduction

Biocatalysis has become increasingly powerful for the efficient synthesis of optically pure pharmaceuticals, agrochemicals, food ingredients, nutraceuticals and fragrances (Borelli and Trono, 2015; Forti et al, 2015; Narancic et al., 2015; Min et al. 2016; Truppo, 2017; Fernández-Lucas et al., 2017). A key challenge in biocatalysis is to overcome the poor solubility in aqueous media for many organic molecules of interest (Makshakova et al., 2012; Karbalaei-Heidari et al., 2013). While biocatalysts are often limited in terms of their stability under industrial process conditions, directed evolution and rational enzyme engineering can potentially be used to address many of these issues, which include poor stability at extremes of pH, temperature and under oxidative stress (Widersten, 2014; Denard et al., 2015; Morris et al., 2016).

Many syntheses involve organic substrates or products that are poorly soluble in water, or are sensitive to aqueous degradation, resulting in a need for enzymes that operate efficiently in organic media, aqueous-organic mixtures or aqueous two-phase systems (Börner et al. 2017; Kong et al, 2017). Many enzymes can function in both aqueous and near-anhydrous organic solvents (Vossenberg et al. 2013), and in some cases organic solvents are found to improve enzyme stability or even alter their enantioselectivity (Klibanov, 2001; Lotti et al., 2015). The impact of solvents on protein structure and dynamics (Duboué-Dijon et al. 2017) have been characterised extensively in near-anhydrous systems where the enzymes are typically freeze-dried or chemically cross-linked prior to addition to organic solvents (). However, most enzymes are rapidly inactivated even at low concentrations of organic co-solvents, and yet the mechanisms of activity loss remain unclear (Stepankova et al., 2013). Activity loss is typically solvent concentration dependent, but varies with organic solvent, potentially depending on solvent hydrophobicity (Paggiola et al. 2014).

Enzymes typically have lower reaction rates in organic media relative to those in aqueous solutions, and yet they are often found to retain their overall native structure (Sinha and Khare, 2014). However, direct structural investigations of enzymes that are either suspended or solubilized into organic media are difficult, and so in the past, activity assays have been relied upon to give an indirect probe of protein structure and function. As a result, the mechanisms leading to partial inactivation of enzymes in organic solvents are still not clear, or generalizable to a wide range of enzymes such that they elucidate the relative roles of protein-solvent interactions, surface or active-site dehydration, structural denaturation, partial unfolding, subunit or cofactor dissociation, and enzyme aggregation (Ahmad et al., 2008, 2012; Mukherjee and Gupta, 2017).

To take advantage of the benefits of organic solvents in biocatalysis, many efforts have been made to enhance enzyme activity and stability in aqueous-organic mixtures, particularly using enzyme immobilisation (Konieczny et al., 2014; Diener et al., 2016; Gupta, 2016) and directed evolution (Illanes et al. 2012; Fei et al. 2015)). These studies have provided useful insights into the impact of organic solvents on enzyme structure and activity. However, novel rational or computational design of organic solvent tolerant enzymes, and their directed evolution, could be significantly improved if the mechanisms by which organic solvents cause enzyme inactivation were better understood.

*E. coli* transketolase (TK) (EC 2.2.1.1) is a homodimeric enzyme in which each 78 kDa subunit comprises three domains, and the apo-TK homodimer binds two Ca^2+^ or Mg^2+^ ions, and two thiamine diphosphate (TPP) cofactors to form the active holo-TK enzyme (Kochetov et al., 1975; Sundström et al., 1992). It catalyses the stereoselective transfer of a two-carbon ketol group to an aldose sugar producing a new asymmetric carbon-carbon bond, at two steps within the reductive pentose phosphate pathway (Turner et al., 2000). The reaction is rendered irreversible *in vitro* by the release of carbon dioxide, when using β-hydroxypyruvate as the ketol donor. These features make the enzyme attractive for the biocatalytic synthesis of complex carbohydrates and their analogues (Liqvist *et al*, 1992; Hecquet *et al*, 1994, 1996), as applied in the industrial synthesis of xylulose-5-phosphate (Woodley, et al., 1996; Shaeri, et al. 2008).

The synthetic potential of TK has already been significantly improved by successive rounds of directed evolution with smart libraries (; Affaticati et al., 2016; Yu et al. 2017), to enhance activity towards polar (Hibbert et al., 2007; Subrizi et al., 2016), and increasingly aliphatic (Hibbert et al., 2008; Cazares et al., 2010: Strafford et al., 2012), and heterocyclic substrates (Cazares et al. 2010; Galman et al., 2010), to enhance and reverse enantioselectivity (Smith et al., 2008). *De novo* pathways containing improved TK variants have also been designed for the synthesis of optically pure high-value amino-diols (Rios-Solis et al., 2011, 2015). Most recently the “substrate-walk” by directed evolution was continued to introduce new activity towards substituted benzaldehyde substrates (Payongsri et al., 2015). However, many new benzaldehydes were too insoluble to test their activity, which has increased the need to explore the use of TK variants in the presence of organic co-solvents.

While we have previously characterised the activity, thermostability and aggregation of transketolase at a wide range of pH, temperature and chemical denaturant concentrations (Martinez-Torres et al., 2007; Jahromi et al., 2011), its stability in a range of organic co-solvents has not been previously determined. TK unfolds irreversibly with urea, heat, or at low pH, and then aggregates, except in urea. Folded yet inactive states form during early urea-unfolding, and at high pH, in which the cofactors remain bound. The enzyme then partially unfolds at higher urea prior to dissociation of the monomers.

Several differences in the protein conformation of the holo and apo TK has been previously observed, for example the secondary and tertiary structure of holo-TK remained constant from pH 5 to 11, whereas the apo-TK secondary structure content increased with pH (Jahromi et al., 2011). Additionally, effects of varying cofactor concentrations on TK activity in holo-TK and apo-TK have been observed, which the latter showed inactivity in the lysate wild-type TK (Miller et al., 2007). Recently, a mutation to proline in one of the two TK cofactor-binding loops was found to stabilise TK to aggregation at elevated temperatures (Morris et al., 2016); and based on this, flexible loops within the structure were engineered and found improvement in thermostability (Yu et al. 2017). Thus, one or more of several potential mechanisms could influence TK stability and activity in solvents, including: i) global unfolding; ii) aggregation; iii) partial unfolding of local structure or domains; iv) cofactor dissociation; and v) modulating the accessibility of water molecules or substrates into the active site.

Here we have characterised the impact of different solvents used in the pharmaceutical industry, including acetonitrile (MeCN), *n*-butanol (nBuOH), ethyl acetate (EToAc), isopropanol (iPrOH), and tetrahydrofuran (THF), on both apo-TK and holo-TK activity and conformation, and correlated this to the physico-chemical properties of the solvents. We also determined the extent to which global and local protein unfolding, and aggregation play a role in solvent-induced enzyme inactivation. This work has provided useful insights that will guide the future engineering of TK and other similarly complex enzymes.

## 2. Materials and methods

All chemicals and solvents were obtained from Sigma-Aldrich (Gillingham, UK) unless noted otherwise.

### 2.1. pQR791 purification

N-terminally His_6_-tagged wild-type *E. coli* transketolase was expressed from *E. coli* XL10-Gold (Stratagene, La Jolla, CA) containing the engineered plasmid pQR791, purified as described previously (Martinez-Torres et al., 2007), dialysed at 4 °C for 24 h against 25 mM Tris-HCl, pH 7.0, and stored at 4 °C for a maximum of two weeks without loss of activity, and with no precipitation visible. Protein concentration was determined by absorbance at 280 nm, assuming a monomeric molecular weight (MW) of 72260.82 g mol^-1^ and an extinction coefficient (ε) of 93905 L mol^-1^ cm^-1^ (Martinez-Torres et al., 2007).

### 2.2. Residual activities after incubation with organic solvents

Samples were divided into two groups. The first determined the impact of solvents on holo-TK. 40 μL of 2.6 mg mL^-1^ apo-TK (in 125 mM Tris-HCl, pH 7.0) was mixed with 5 µL of a 20x cofactor stock (48 mM TPP, 180 mM MgCl_2)_. This was incubated for 20 min at 25 °C with 1000 rpm shaking, before adding 55 µL of a range of solvents at different concentrations in water to yield 1 mg mL^-1^ holo-TK solutions (2.4 mM TPP, 9 mM MgCl_2_) in 50 mM Tris-HCl, pH 7.0. These solutions were then incubated for 3 h at 25 °C 1000 rpm. For the second group, apo-TK solutions received the same treatment as the described above for holo-TK, expected that 5 µL of water were added instead of the 5 µL of the 20x cofactor stock solution.

All residual activities were determined by addition of 15 µL enzyme solution samples to 285 µL of substrate stock (in 1x cofactors and 50 mM Tris-HCl, pH 7.0), giving final concentrations of 50 mM HPA, 50 mM glycolaldehyde, 2.4 mM TPP, and 9 mM MgCl_2_ in 50 mM Tris-HCl, pH 7.0, and 20-fold dilution of the original solvent. At regular intervals during a period of 90 min, aliquots of 20 µL were diluted 1:10 with 180 µL 0.1% (v/v) TFA, to stop the reactions and then transferred into 96 micro-well plates for measuring of products by HPLC as described previously (Hibbert et al, 2007). Samples were loaded on a 300mm Aminex HPX-87H column (Bio-Rad Laboratories) maintained at 60 °C, and analysed with an isocratic flow of 0.1% (v/v) TFA in water at 0.6 mL min^−1^.

### 2.3. Secondary structure monitored by circular dichroism (CD)

CD spectra (190-300 nm) were recorded on an AVIV 202 SF spectrometer (AVIV Associates, Lakewood, NJ) at 25 °C using a 1 mm path length quartz precision cell cuvette. Samples were prepared as above at 0.5 mg mL^-1^ transketolase both with and without 2.5 mM MgCl_2_ and 0.25 mM TPP for holo-TK and apo-TK respectively, in 25 mM Tris-HCl, pH 7.0, and with the required % (v/v) solvent. The lower cofactor concentrations are sufficient for >90% saturation of the TK (Miller, 2007), while minimising the dynode voltage in CD spectra due to absorbance by the cofactors. CD spectra were recorded at 0.5 nm intervals and averaged for 4 seconds at each wavelength. Spectra were recorded at 30-45 min intervals for up to 15 h after the addition of solvents. Spectra for 25 mM Tris-HCl, pH 7.0 buffer in the respective % (v/v) solvent was subtracted from each recording.

### 2.4. Intrinsic fluorescence intensities

Holo- and apo-TK samples were prepared as above at 0.1 mg mL^-1^ either with or without 5 mM MgCl_2_ and 0.5 mM TPP, in 25 mM Tris-HCl, pH 7.0, and with the required % (v/v) solvent. Samples were incubated at 25 °C in UV transparent 96-well microplates (Costar, Corning Incorporated, NY, USA). Fluorescence intensity was measured every 30 min for 6 h from below the plate at 340 nm emission and 280 nm excitation using a FLUOstar microplate reader (BMG Labtechnologies Ltd., Aylesbury, UK). Different gain settings were used for the apo-TK and holo-TK experiments. Due to the interference with measurements by an incompatibility of EToAc with the sample plates that appears after long incubations, those incubations were carried out in glass vials before transferring samples to UV transparent 96-well microplates immediately prior to measurements.

### 2.5. Dynamic light scattering (DLS)

The particle size distributions of transketolase in the presence and absence of solvent was measured at 25 °C with a Zetasizer Nano S (Malvern Instruments Ltd., UK). Holo-TK at 0.1 mg mL^−1^ was prepared as above in 25 mM Tris-HCl, pH 7.0, with 0.5 mM TPP and 5 mM MgCl_2_ and the required % (v/v) solvent. Samples were incubated for 1, 2 and 3 h prior to data acquisition. Data were acquired in triplicate, with a 1 cm path length low volume disposable sizing cuvette. Control samples of the respective buffered solvents, with or without cofactors was subtracted from each recording. The hydrodynamic diameters of each sample were calculated from the averaged-measurements using the Zetasizer Nano Series software V.4.20 (Malvern Instruments Ltd., Worcestershire, UK).

## 3. Results and discussions

### 3.1 Transketolase activity in organic co-solvents

To understand the effect of polar organic co-solvents upon TK stability and activity, as well as the influence of cofactors, both apo-TK and holo-TK were incubated with five selected co-solvents at a range of concentrations from 0-30% (v/v). In the first case, apo-TK was incubated with the organic solvents for 3 h and then incubated with the cofactor to form the holo-TK before diluting and adding the substrates. In the second case, to test holo-TK, the apo- TK was first incubated with cofactors to form the holo-TK, and then incubated with organic solvents for 3 h before diluting and adding the substrates. In both cases, the solvent incubations were shaken at 1000 rpm to ensure that the least miscible solvents, ethyl acetate (EToAc), and *n*-butanol (nBuOH) were either fully dissolved or formed into emulsions at the higher concentrations (biphasic systems were only formed for EToAc and nBuOH at concentrations of 8.3 and 7.3% v/v respectively).

Cofactor-free effects were studied to give additional insights to TK activity (Miller et al. 2007). The experiments were done as described above using four solvents: THF, iPrOH, EToAc and MeCN; the results are shown in Fig. 2, where it can be seen that the enzyme activity is 286-fold less active than the holo-TK at co-solvent-free system and decreased gradually. With 25% (v/v) MeCN, complete inactivation was achieved and at 30% (v/v) of the rest of the co-solvents except EToAc remained slightly active.

**Fig. 1.**
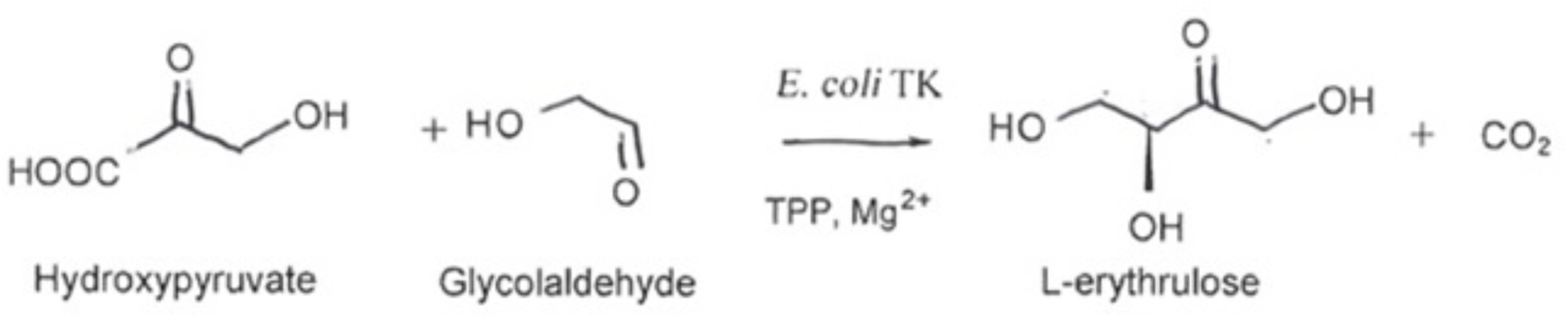
Reaction of transketolase

**Fig. 2.**
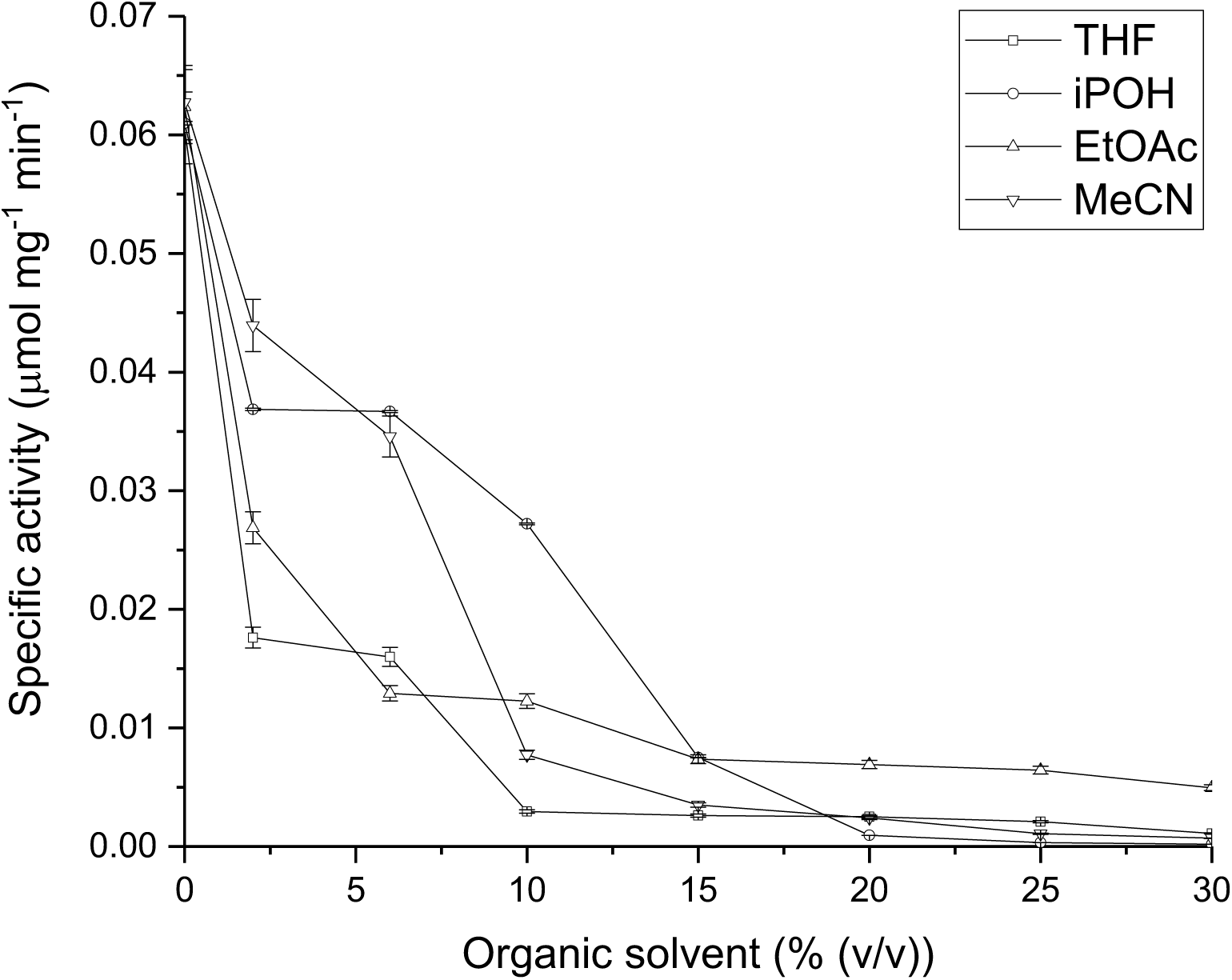
Retention of transketolase catalytic activity in cofactor-free systems with increasing concentrations of organic solvents. Apo-TK was compared for its stability to incubation with (□) THF, (°) isopropanol, (Δ) ethyl acetate and (▽) acetonitrile, in 50 mM Tris-HCl, pH 7.0 (plus 4.8 mM TPP, 18 mM MgCl_2_ for holo-TK), for 3 h at 25°C and 1000 rpm shaking. Enzyme activity was measured after a 20-fold dilution of the solvents, at 50mM HPA and 50 mM GA in 2.4 mM TPP, 9 mM MgCl_2_, 50 mM Tris-HCl, pH 7.0. Error bars represent one standard deviation about the mean (n = 3).

The results shown in Fig. 3 compare the effect of increasing the final solvent concentrations, upon the activity retained after 3 h, for apo-TK and holo-TK.

**Fig. 3.**
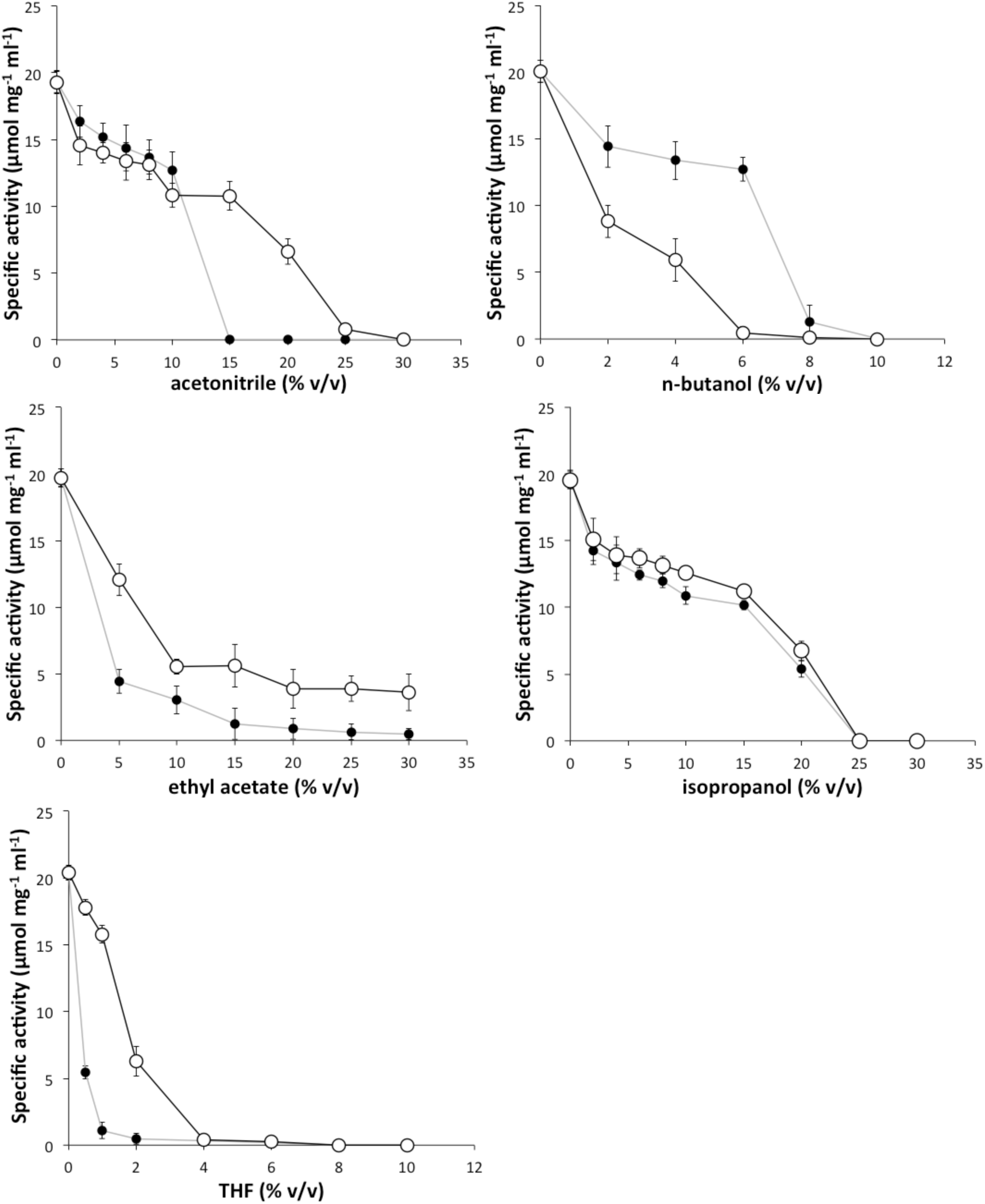
Retention of transketolase catalytic activity after pre-incubation with increasing concentrations of organic solvents. (○) Apo-TK and (•) holo-TK were compared for their stability to incubation with (a) acetonitrile (b) *n*-butanol (c) ethyl acetate (d) isopropanol (e) THF, in 50 mM Tris-HCl, pH 7.0 (plus 4.8 mM TPP, 18 mM MgCl_2_ for holo-TK), for 3 h at 25°C and 1000 rpm shaking. Enzyme activity was measured after a 20-fold dilution of the solvents, at 50mM HPA and 50 mM GA in 2.4 mM TPP, 9 mM MgCl_2_, 50 mM Tris-HCl, pH 7.0. Error bars represent one standard deviation about the mean (n = 3).

**Fig. 4.**
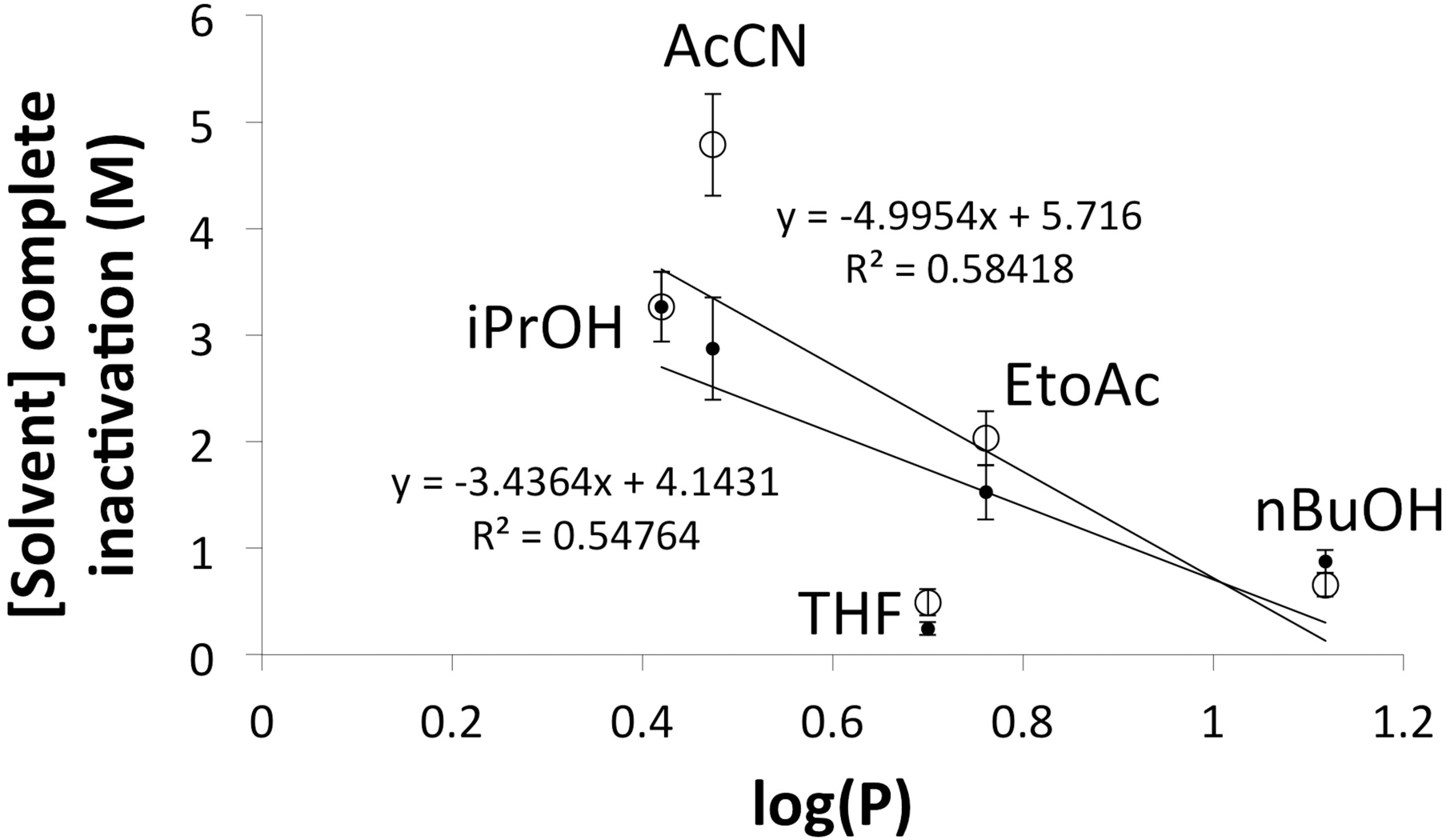
Correlations of solvent log(P) with their TK inactivation potency. Log(P) correlates well with the molar concentration required to inactivate either (**□**) holo-TK or (♦) apo-TK. Error bars represent one standard deviation about the mean (n = 3).

In nearly all cases, the increased concentration of co-solvents eventually reduced the remaining activity to zero. Apo-TK was more stable than holo-TK to three of the solvents tested, but less stable than holo-TK in nBuOH, and essentially of the same stability as holo-TK in iso-propanol (iPrOH). For example, acetonitrile (MeCN) half-inactivated apo-TK and holo-TK at 20% (v/v) and 12% (v/v), respectively. Similarly, EToAc half-inactivated holo-TK at approximately 3% (v/v), and at 6% (v/v) EToAc for apo-TK. Nevertheless, apo-TK retained 18% residual activity at 30% (v/v) EToAc, whereas holo-TK was already completely inactivated by 15% (v/v). The sharp decrease in activity up to 10% EToAc and the apparent retention of activity above it for both apo and holo-TK potentially resulted from having reached the 8.3% (v/v) solubility of EToAc in water (Altshuller and Everson, 1953). Increasing the concentration at above 8.3% (v/v), formed emulsions which affected the deactivation profile of apo-TK and holo-TK. Holo-TK was completely inactivated by 15% (v/v), but not apo-TK, even at 30% (v/v). Inactivation of enzymes in emulsions is a complex process, which reflects an expected increase in the surface area of the solvent-water interface, which in turn promotes transfer of solvent molecules to the protein surface.

In iPrOH, both apo-TK and holo-TK behaved similarly with a gradual drop in activity until there was no activity in either case at 25% (v/v) iPrOH. Tetrahydrofuran (THF) was the most effective at deactivation, with holo-TK completely inactivated at just 2% (v/v) THF and apo-TK at 4% (v/v) THF. By contrast, apo-TK was less stable to nBuOH than holo-TK with concentrations of 2% (v/v) and 7% (v/v) respectively giving rise to 50% activity. For both apo-TK and holo-TK in nBuOH the residual activity tended to zero after reaching the 7.3% (v/v) solubility limit for nBuOH. This sharp decrease in stability for apo-TK was probably due to the solvent-water interface effect.

The generally lower tolerance of holo-TK to co-solvents compared to apo-TK is counterintuitive given that holo-TK is thermodynamically more stable than apo-TK (Martinez-Torres et al., 2007; Esakova et al., 2005). The dependence on solvent concentration is also clearly not simple. For example, MeCN, nBuOH and iPrOH displayed an initial lag or plateau at low solvent concentrations, where the activity decreases only slightly, before reaching a critical concentration at which deactivation occurs more abruptly. Such sigmoidal profiles can indicate a cooperative transition for structural unfolding or even the dissociation of protein dimers into monomers. By contrast, THF and EToAc titrated out the activity rapidly with increasing concentration of solvent, suggesting either an isotherm for solvent binding to the protein, or that any cooperative transition was already well underway at the lowest concentration of solvent tested.

As apo-TK is already known to be less thermodynamically stable than holo-TK, and yet was found typically to be more solvent tolerant, TK inactivation by solvent does not appear to relate simply to the global conformational stability (or global denaturation) of the protein. Further biophysical characterisations were undertaken to elucidate whether inactivation was due to global protein denaturation, local protein unfolding, protein aggregation, or some other effect on the protein structure that is not directly detected, such as active-site binding. Before that we determined any relationships between the properties of the solvents and their potency for enzyme inactivation.

### 3.2 Correlation of TK activity to calculated organic solvent properties

Although all the organic-solvents in these experiments were polar, they showed considerable variability in their impact on retained TK activity, including their critical concentrations for inactivation, the sharpness of their deactivation curves, and also in their relative impacts upon apo-TK and holo-TK. The effectiveness of a polar solvent for inactivation of proteins might be expected to depend upon the polarity or hydrophobicity of the solvent, and also their ability to form hydrogen bonds in place of water.

The polar solvents can be categorized as aprotic (MeCN, EToAc and THF), or protic (nBuOH and iPrOH). This simple categorization had no obvious correlation to the concentrations of co-solvents required for complete enzyme inactivation. The characteristics of each solvent used in terms of various theoretically calculated and experimentally determined physicochemical properties, is shown in Table 2, along with the Pearson’s R^2^ values for linear correlations between each solvent property, and the molar solvent concentrations required for complete inactivation of holo-TK. Log(P) gave the best correlation 4for both apo-TK (R^2^=0.58) and holo-TK (R^2^=0.55) (Supplementary Figure S1), although those correlation values are still poor. Other solvent properties correlated even less with the molar holo-TK inactivation concentration, such as the molecular weight (R^2^=0.4), molecular volume (R^2^=0.4), topological polar surface area (R^2^=0.3), number of potential hydrogen bonds (R^2^=0.1), dipole moment (R^2^=0.2), and the dielectric constant of the co-solvent (R^2^=0.4). Although the Log(P) correlation was poor, it showed the highest R^2^ which was consistent with previous observations that correlated hydrophobicity, and not the dielectric constant, to loss of enzyme activity and stability (Esakova et al., 2005; Miller et al., 2007; Galman et al., 2010; Jahromi et al., 2011; Strafford et al., 2012). However, various mechanisms have been used to explain such previous correlations, and so we sought to further characterise the solvent-induced inactivation of TK, as summarised in Table 3 and described below.

**Table 1.** Pharmaceutical applications of the chosen polar co-solvents

| Co-solvent | Acronym | Common industrial applications | Reference in open literature |
| --- | --- | --- | --- |
| 2-propanol | iPrOH | Substitute for ethanol in extracts and tinctures | Burlage and Hawkins, 1946; Schaller and Triebig, 2012 |
| n-butanol | nBuOH | Reacting intermediate in the synthesis of other chemicals as a solvent and as an ingredient in formulated products such as antibiotics, hormones, and vitamins | DOW, 2013 |
| acetonitrile | MeCN | Reacting intermediate in the synthesis of pharmaceutical products, such as: Vitamin B1; sulfa pyrimidine, antibiotics | McConvey, et al., 2012 |
| ethyl acetate | EToAc | Extraction agent in the production of pharmaceuticals | Berton et al., 2017 |
| tetrahydrofuran | THF | Widely used as a solvent, adhesive, and for the synthesis of chemicals such as butyrolactone, succinic acid | Yao et al., 2012 |

**Table 2.**
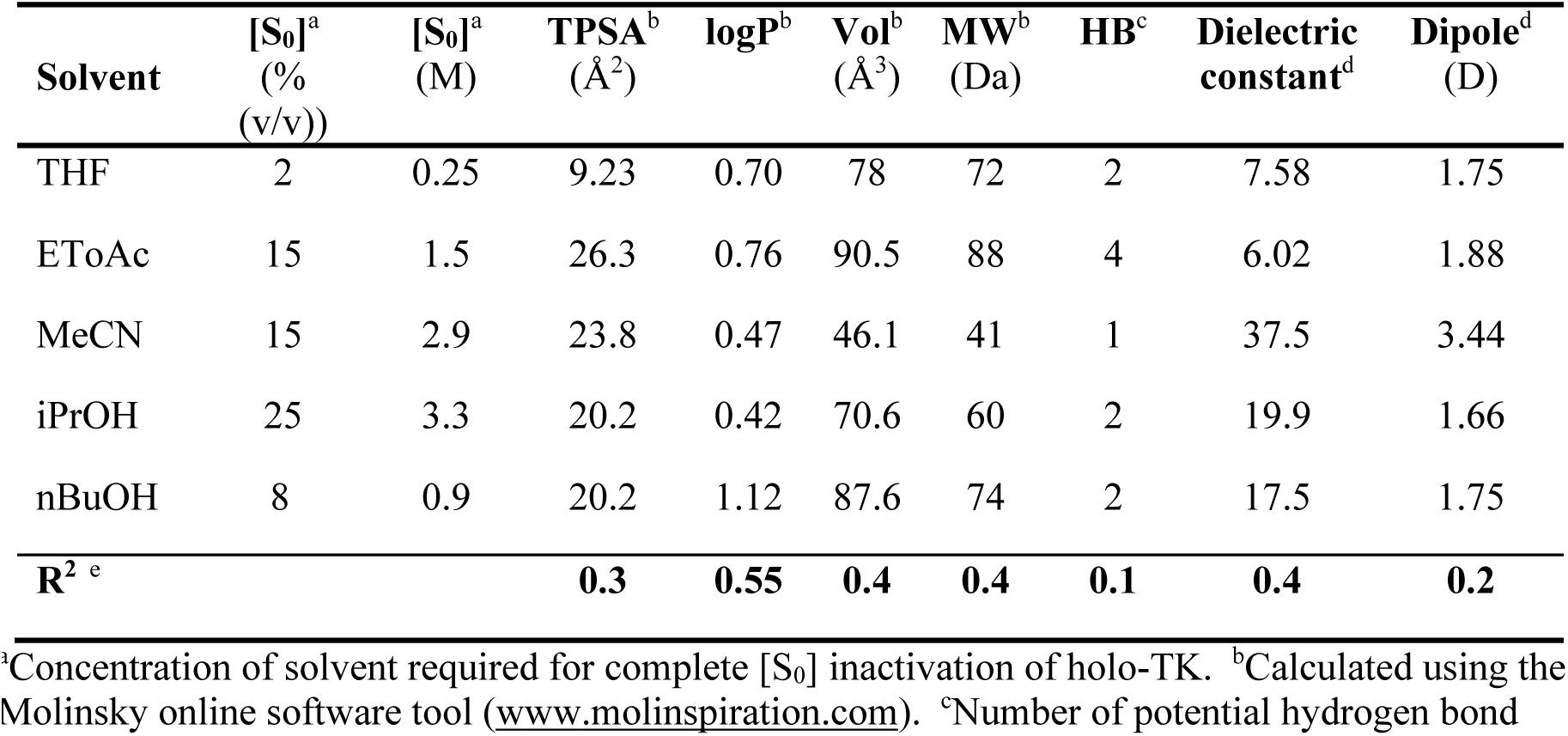
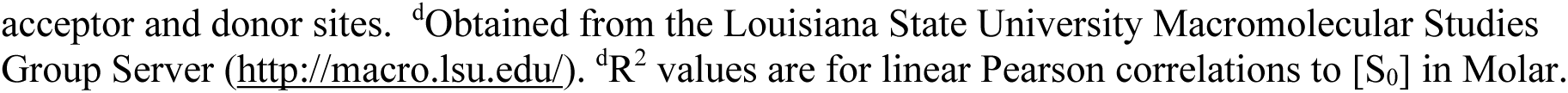
Correlation of holo-TK inactivation to physicochemical properties of the polar organic solvents.

| Solvent | [S<sub>0</sub>]<sup>a</sup><br>(%<br>(v/v)) | [S<sub>0</sub>]<sup>a</sup><br>(M) | TPSA<sup>b</sup><br>(Å<sup>2</sup>) | logP<sup>b</sup> | Vol<sup>b</sup><br>(Å<sup>3</sup>) | MW<sup>b</sup><br>(Da) | HB<sup>c</sup> | Dielectric<br>constant<sup>d</sup> | Dipole<sup>d</sup><br>(D) |
| --- | --- | --- | --- | --- | --- | --- | --- | --- | --- |
| THF | 2 | 0.25 | 9.23 | 0.70 | 78 | 72 | 2 | 7.58 | 1.75 |
| EToAc | 15 | 1.5 | 26.3 | 0.76 | 90.5 | 88 | 4 | 6.02 | 1.88 |
| MeCN | 15 | 2.9 | 23.8 | 0.47 | 46.1 | 41 | 1 | 37.5 | 3.44 |
| iPrOH | 25 | 3.3 | 20.2 | 0.42 | 70.6 | 60 | 2 | 19.9 | 1.66 |
| nBuOH | 8 | 0.9 | 20.2 | 1.12 | 87.6 | 74 | 2 | 17.5 | 1.75 |
| **R<sup>2</sup> <sup>e</sup>** |  |  | **0.3** | **0.55** | **0.4** | **0.4** | **0.1** | **0.4** | **0.2** |

<sup>a</sup>Concentration of solvent required for complete [S<sub>0</sub>] inactivation of holo-TK. <sup>b</sup>Calculated using the Molinsky online software tool (www.molinspiration.com). <sup>c</sup>Number of potential hydrogen bond
acceptor and donor sites. <sup>d</sup>Obtained from the Louisiana State University Macromolecular Studies Group Server (http://macro.lsu.edu/). <sup>d</sup>R<sup>2</sup> values are for linear Pearson correlations to [S<sub>0</sub>] in Molar.

**Table 3.** Effects of polar organic co-solvents on TK structure and activity after 3 h at 25 °C.

| Solvent | [S]<sup>a</sup><br>(%(v/v)) | Activity at 3 h<br>(%) |  | CD<sup>b</sup> % Native<br>Holo TK<br>(±2%) |  | CD Initial rate<br>unfolding<sup>c</sup><br>(% hr<sup>-1</sup>) |  | FLI transition<sup>d</sup><br>(%(v/v)) | DLS |
| --- | --- | --- | --- | --- | --- | --- | --- | --- | --- |
|  |  | Holo | Apo | 0 h | 3 h | Holo | Apo | Holo | Holo |
| None | 0 | 100 | 100 | 100 | 101 | 0 | 0 | n/a | 7 nm |
| THF | 2 | 0 | 32 | 78 | 74 | 1.7 | 1.7 | none | 9 nm |
| EToAc | 10 | 15 | 31 | 99 | 98 | 1.2 | 1.4 | none | 8 nm |
| MeCN | 20 | 0 | 35 | 90 | 54<sup>f</sup> | 16.2 | 1.1 | >15% | >1000 nm |
| iPrOH | 20 | 30 | 35 | 100 | 95 | 1.4 | 1.0 | >22% | 15 nm |
| nBuOH | 8 | 10 | 0 | 98 | 98 | 0.8 | 2.1 | 7-8% | > 1000 nm |

<sup>a</sup>Concentration of solvent added. <sup>b</sup>CD % Native based on signal at 222 nm, except EoAc which is based on signal at 223 nm to avoid a high dynode voltage at <222 nm. <sup>c</sup>Initial rate of unfolding was determined by converting mean residue ellipticity into % loss of native structure, and by subtracting the baseline rate determined in the absence of solvent. <sup>d</sup>Fluorescence intensity. <sup>e</sup>Time dependence of fluorescence intensity (approximate time taken to reach equilibrium). <sup>f</sup>Aggregated after 4.5h.

### 3.3 Secondary structure of apo-TK and holo-TK in polar co-solvents

To determine the impact of co-solvents on the secondary structure content of TK, and also whether the enzyme was unfolding or aggregating, far-UV circular dichroism (CD) spectra were obtained at approx. 30-45 min intervals for between 3 and 22 h, for both apo-TK and holo-TK, using each volume fraction of solvent that resulted in ≈70% inactivation of the apo-TK enzyme (specific activity of 6 μmol mg^-1^ min^-1^), except for nBuOH which was increased to ensure holo-TK was also inactivated.

The time dependencies of the mean residue ellipticity at 222 nm for apo-TK and holo-TK in each co-solvent, and also without co-solvent, are shown in Fig. 5A, and the initial rates are shown in Table 3. The far-UV CD spectra of holo-TK, after incubation with co-solvents for 3 h, are also shown in Fig. 5B. It can be seen in Fig. 5A that the control samples of apo-TK and holo-TK with no added co-solvent, each gave no statistically significant change in secondary structure content over the course of at least 4 h. By contrast, relative to the control samples, the co-solvents induced a small initial unfolding rate of between 1.0 and 2.1% hr^-1^ loss of native structure for apo-TK, and between 0.8 and 1.7% hr^-1^ for holo-TK, except for MeCN with holo-TK which unfolded more rapidly at 16% hr^-1^. Apo-TK and holo-TK had similar unfolding rates in any given solvent, except for in MeCN where holo-TK unfolded 15-fold faster than apo-TK, and conversely for nBuOH where apo-TK unfolded 2.7-fold more rapidly than holo-TK. MeCN with holo-TK was the only sample for which aggregates were detected by CD as indicated after 4.5 h of unfolding, by sudden and simultaneous increases in the dynode voltage and mean residue ellipticity.

**Fig. 5A.**
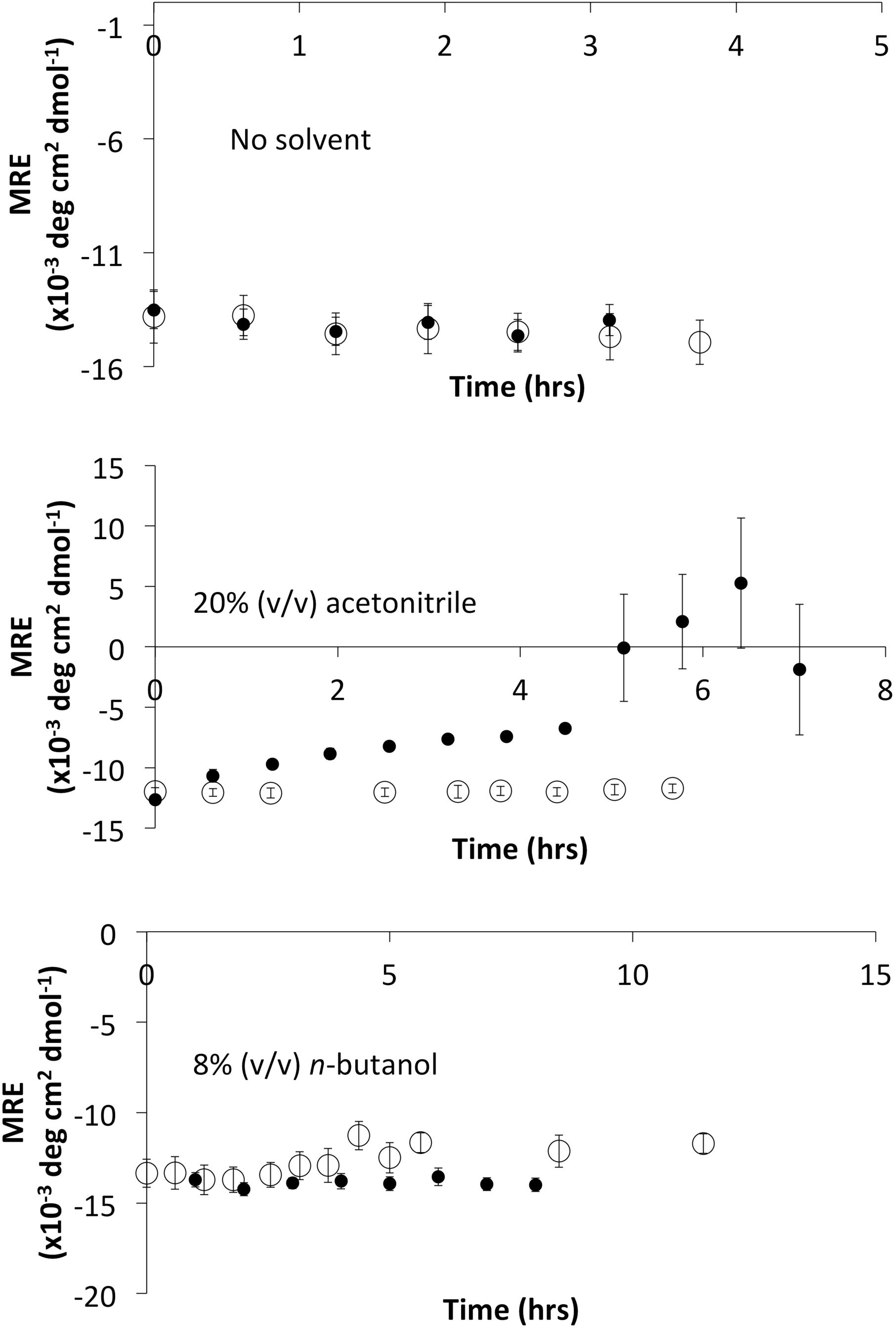
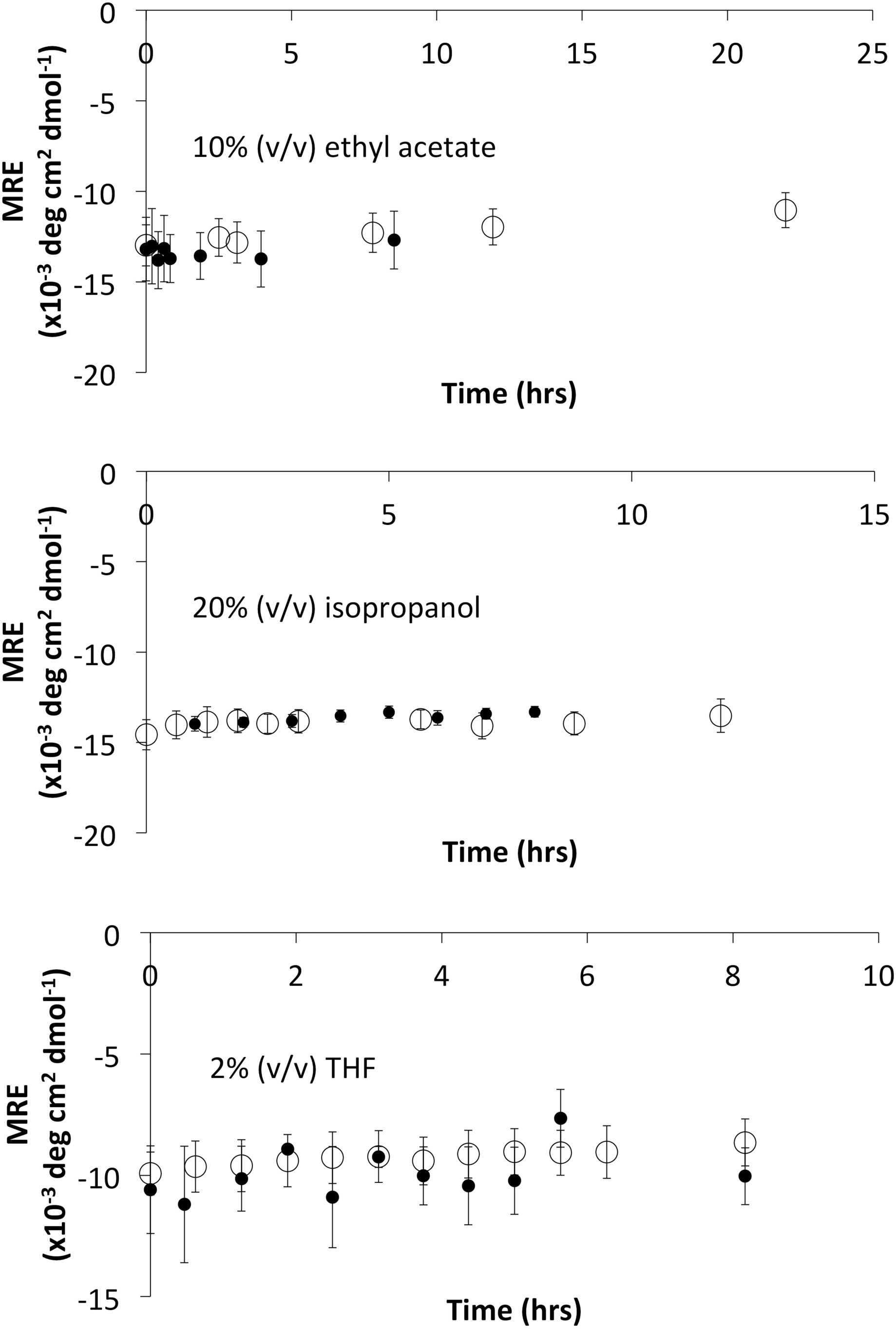
Time-dependence of circular dichroism mean residue ellipticity at 222 nm, after mixing with co-solvents. (**○**) Apo-TK and (•) holo-TK at 0.5 mg mL^−1^ in 25 mM Tris-HCl, pH 7.0, were incubated with no solvent, 20% (v/v) acetonitrile, 8% (v/v) *n*-butanol, 10% (v/v) ethyl acetate, 20% (v/v) isopropanol, or 2% (v/v) THF. Holo-TK also contained 2.5 mM MgCl_2_, 0.25 mM TPP. Error bars represent one standard deviation about the mean (n = 3).

**Fig. 5B.**
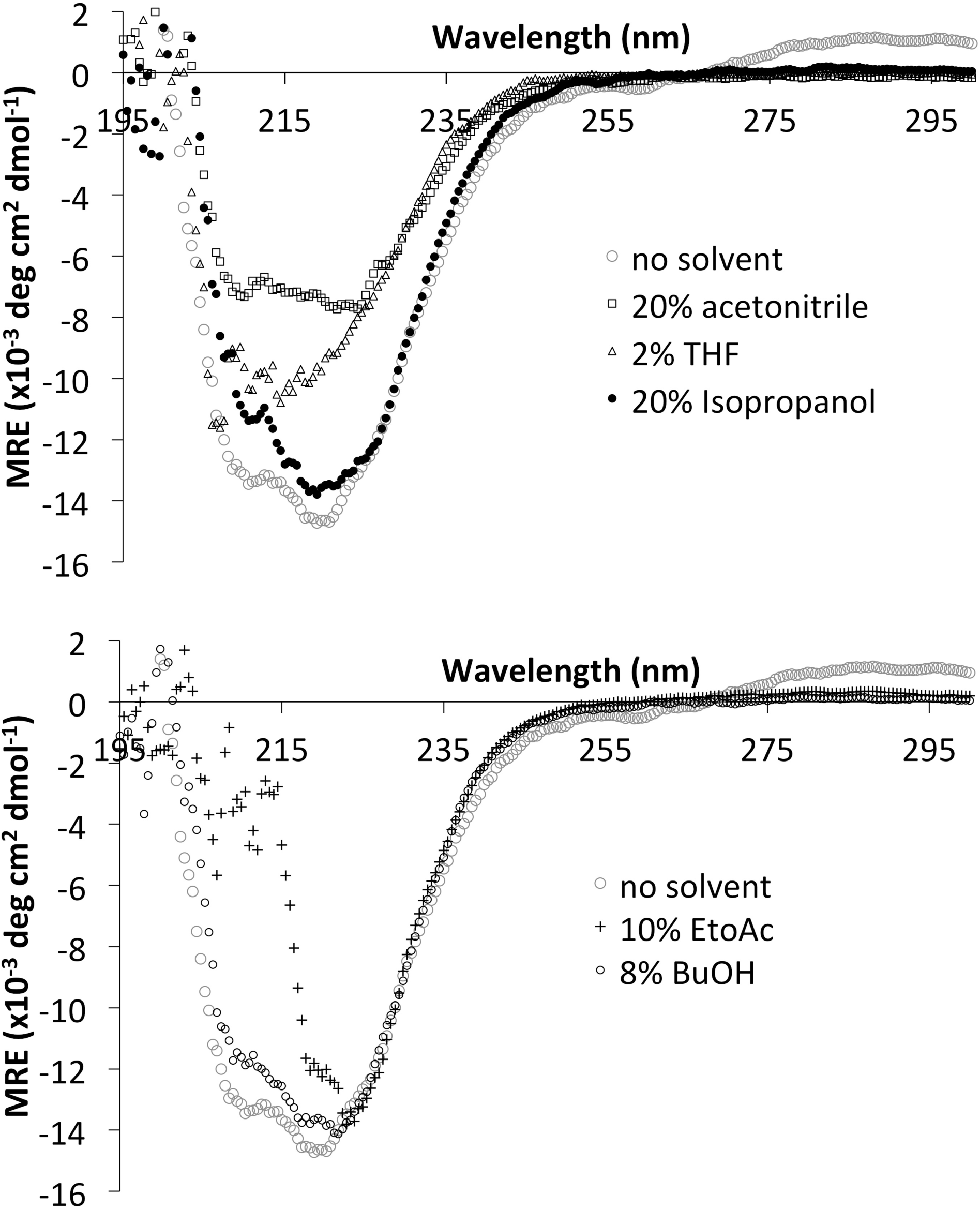
Circular dichroism spectra of holo-TK after incubation with organic solvents. Holo-TK (0.5 mg mL^−1^) in 25 mM Tris-HCl, pH 7.0, 2.5 mM MgCl_2_, 0.25 mM TPP, and the presence of (**○**) no solvent; (**□**) 20% acetonitrile; (**○**) 8% *n*-butanol (+) 10% ethyl acetate (•) 20% isopropanol (Δ) 2% THF, was incubated for 3 h at 25 °C before full spectra (195–300 nm) were acquired. Error bars represent one standard deviation about the mean (n = 3).

It can be seen from Fig. 5A that all samples, except for THF, began with the same initial secondary structure content, with a mean residue ellipticity of -13500 ±500 deg cm^2^ dmol^-1^. THF by contrast appeared to begin with only 78% of the native structure (−10590 deg cm^2^ dmol^-1^), yet this only decreased to 74% after 3 h. The far-UV spectrum taken after 3 h in THF (Fig. 5B), and at all earlier time-points (not shown), indicated that the smaller ellipticity at 222 nm was due to distortion from the spectrum expected from a predominantly α-helical protein. This was due to absorbance flattening by THF, that gave a correspondingly increased dynode voltage at 224 nm and below (not shown), to above that acceptable for the instrument, rather than resulting from any actual loss of native structure or conformational change immediately after the addition of THF. EToAc was similarly affected by a high dynode voltage at below 222 nm leading to considerable signal scattering at below 215 nm at all time-points.

From Fig. 6A&B it can be seen that the initial unfolding rates determined by CD were found to correlate well (R^2^ of 0.91 for apo-TK, 0.71 for holo-TK, and 0.73 combined) to the activity retained after 3 h of incubation with the co-solvents, and also to the log(P) of the co-solvents (R^2^ of 0.94 for apo-TK), when excluding the case of MeCN for holo-TK. This indicated that the inactivation of the enzyme by co-solvent for both apo-TK and holo-TK was linked to the slow unfolding observed in most samples. Furthermore, as unfolding of only 0.8-2.1% hr^-1^ native structure led to 70-100% inactivation in apo-TK and holo-TK, then the slow unfolding was most likely to be due to local unfolding of a key element of structure that affects the entire protein population, rather than global unfolding of only 2.1% of the protein population.

**Fig. 6A&B.**
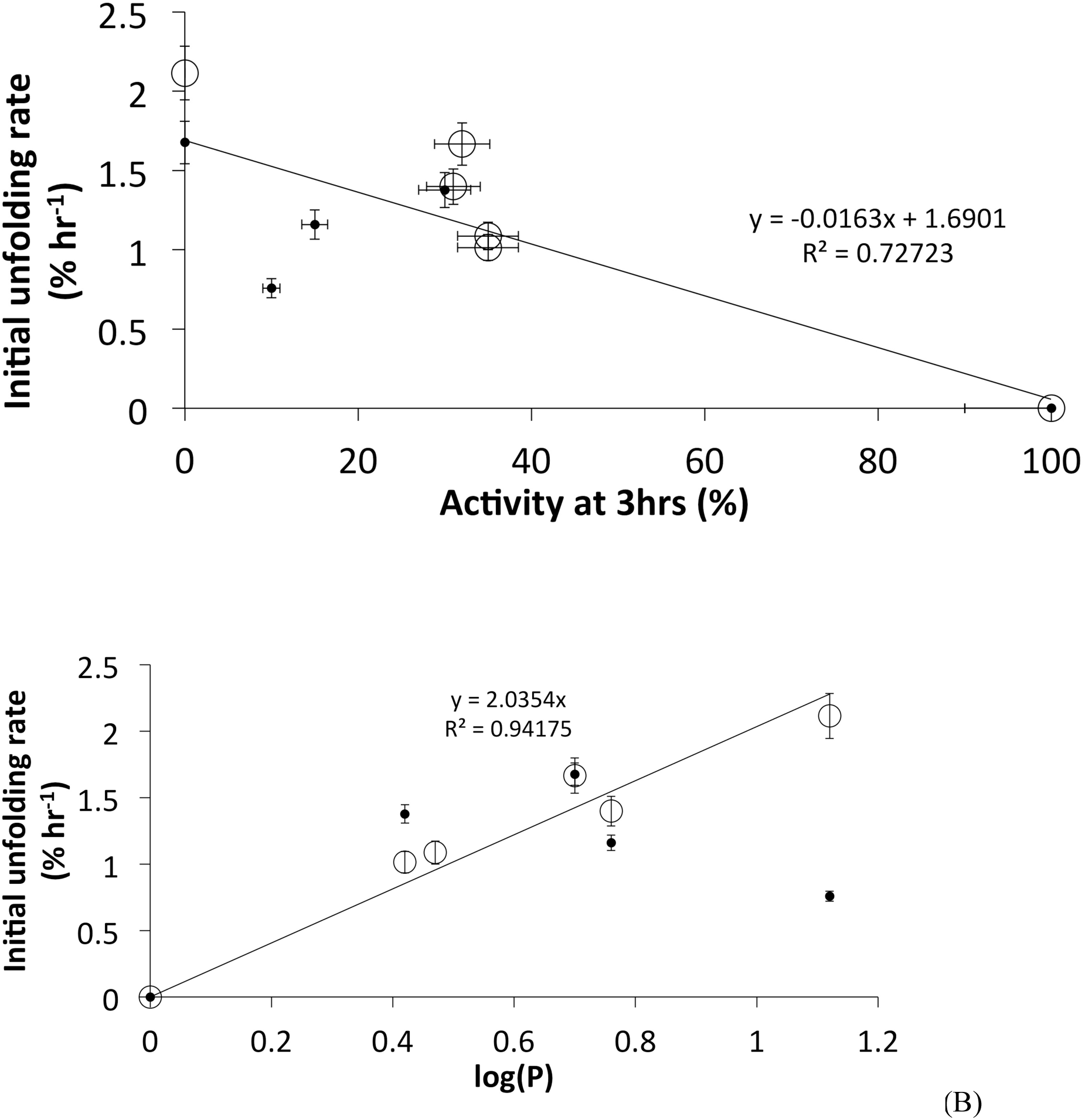
Relationship between the local unfolding rate, retained activity after 3 h, and log(P). Both plots exclude MeCN as this solvent unfolded the protein globally. Apo-TK (**○**) and holo-TK (•).Error bars represent one standard deviation about the mean (n = 3).

For AcCN with holo-TK, the significantly higher rate of unfolding, led to 54% native structure after 3 h, and indicated that AcCN had induced global unfolding, from which aggregation then occurred after 4.5 h. Holo-TK was completely inactive after 3 h, indicating that global unfolding could only account for up to 54% of the inactivation mechanism, and therefore that the local unfolding observed with apo-TK in AcCN, had additionally inactivated holo-TK.

### 3.4 Particle size distributions from dynamic light scattering

Dynamic light scattering (DLS) was used to measure particle size distributions of holo-TK in the presence of the co-solvents (see supplementary Fig. S1). Holo-TK in the absence of organic co-solvent gave a particle size of 7-8 nm as expected theoretically from the native homodimer structure (Littlechild et al., 1995). At 20% (v/v) AcCN and 8% (v/v) nBuOH, holo-TK at 0.1 mg ml^-1^ contained DLS-detectable aggregates of 1000-4000 nm within 1 hour of incubation. At 20% (v/v) iPrOH a very small aggregate peak at 400 nm was observed within 1 hour, but more significantly, the monomer peak had shifted to approximately 15 nm indicating either the formation of a soluble oligomer, a shifted monomer peak due to viscosity effects, the formation of a solvent boundary layer, or otherwise significant protein unfolding. In 10% (v/v) EToAc, only the native peak was detected at 8 nm indicating the retention of the native homodimer, and no aggregation. For 2% (v/v) THF, the native peak increased slightly within 1 hour to 9 nm, indicating a native homodimer with the peak-shift due to altered solvent viscosity, although not inconsistent with partial local unfolding, or the formation of a significant solvent boundary layer associated with the homodimer. No aggregates were detected with THF.

It should be noted that by DLS, the % volume is proportional to the cube of the particle diameter, and hence even very small amounts of aggregates can suppress the detection of the native monomer peak. Therefore, it is very possible that the monomer, or even the 15 nm state observed for iPrOH, was present also for AcCN or nBuOH, but suppressed by the presence of larger (>1000 nm) aggregates. In iPrOH, the smaller (400 nm) aggregate was clearly a very low-populated aggregate which allowed the 15 nm state to be observed. Overall, these results are consistent with the emergence of low levels of aggregates in nBuOH and iPrOH, more significant aggregation in AcCN as detected also by CD, while no aggregation was observed by any method in EToAc or THF. Aggregation itself therefore does not appear to be well correlated to enzyme inactivation, as all solvents led to at least 70% inactivation of holo-TK. Only the significant levels of aggregate observed by CD could potentially contribute to enzyme inactivation for holo-TK, but this only occurred after 4.5 h of incubation, and for one solvent only (AcCN).

### 3.5 Tertiary structure of apo-TK and holo-TK in polar co-solvents

Fluorescence tryptophan spectroscopy has been previously successfully applied to provide tertiary structural information for TK (Esakova et al., 2005; Martinez-Torres et al., 2007; Aucamp et al. 2008; Jahromi et al., 2011). The unfolding of *E. coli* TK can lead to both an increase or decrease in intrinsic fluorescence intensity, as observed previously where urea denaturation gave an initial increase followed by a decrease at higher urea concentrations (Martinez-Torres et al., 2007). For thermal unfolding, the fluorescence intensity of TK increased in a single cooperative transition due to rapid aggregation upon unfolding.

Fig. 7 shows the time-dependence of intrinsic fluorescence intensities, for holo-TK in 5-30% (v/v) co-solvent. EoAc and THF both resulted in steady decreases in fluorescence over time, for all concentrations tested. EToAc at 10% (v/v) led to a 7% decrease in fluorescence, and THF at 5% (v/v) led to a 9% decrease, each after 3 h of incubation. These results are consistent with the local unfolding observed by CD above. By contrast, the fluorescence intensities for AcCN, nBuOH and iPrOH, resulted in lag-phases followed by rapid increases in fluorescence. The rapid increase occurred progressively earlier with increasing co-solvent concentration, and for AcCN at least the lag phase disappeared at the highest concentrations. This lag-phase behaviour is typical of aggregation kinetics (Li et al., 2010), and the fluorescence increases occurred in the same samples for which aggregation was observed by DLS. For 20% (v/v) AcCN, the fluorescence intensity increased over a timescale consistent with the global unfolding and aggregation observed by CD. For AcCN, nBuOH and iPrOH, it remained possible that the fluorescence changes indicative of aggregation, were also convoluted with contributions from the local unfolding events observed by CD.

**Fig. 7.**
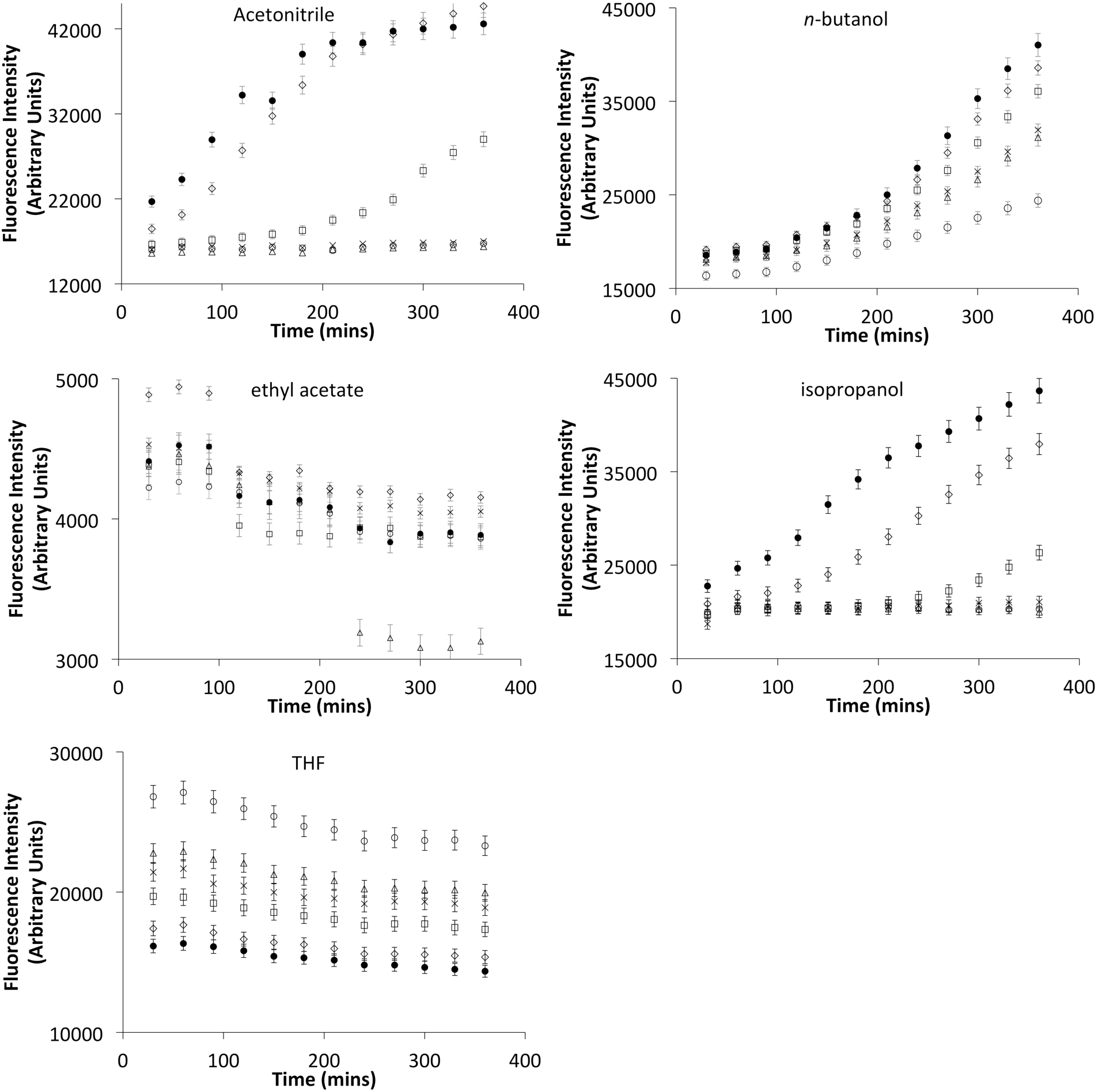
Time dependence of fluorescence intensity at various organic solvent concentrations for holo-TK. Fluorescence intensity was measured at 340 nm with excitation at 280 nm. Holo-TK at 0.1 mg mL^−1^, in 5 mM MgCl_2_, 0.5 mM TPP, 25 mM Tris-HCl, pH 7.0, was measured every 30 min for 6 h at 25 °C in (○) 5%, (Δ) 10%, (x)15%, (□) 20%, (◊) 25% and (•) 30% (a) acetonitrile (b) *n*-butanol (c) ethyl acetate (d) isopropanol (e) THF. Error bars represent one standard deviation about the mean (n = 3).

The effect of increasing co-solvent concentration on intrinsic fluorescence, as measured after 1, 2, and 3 h of incubation, is shown for apo-TK in Fig. 8 and holo-TK in Fig. 9. For apo-TK, the addition of co-solvents resulted in up to a 10% increase in the intrinsic fluorescence after 3 h in AcCN, nBuOH, iPrOH and EToAc (Fig. 7), consistent with local rather than global unfolding. Increases were monotonic for 0-30% (v/v) AcCN, 4-30% (v/v) nBuOH, 10-30 % (v/v) iPrOH, and 0-29% (v/v) for EToAc. Below 10% (v/v) iPrOH there was no change, and at 30% (v/v) EToAc there was a small increase. For THF, the fluorescence intensity decreased exponentially, with most of the change completed over 0-10% (v/v). The lack of obvious transitions for AcCN, EToAc and THF, and yet clear transitions at 0-4% (v/v) nBuOH, and 10% (v/v) iPrOH, were consistent with the inactivation profiles for apo-TK in Fig. 3.

**Fig. 8.**
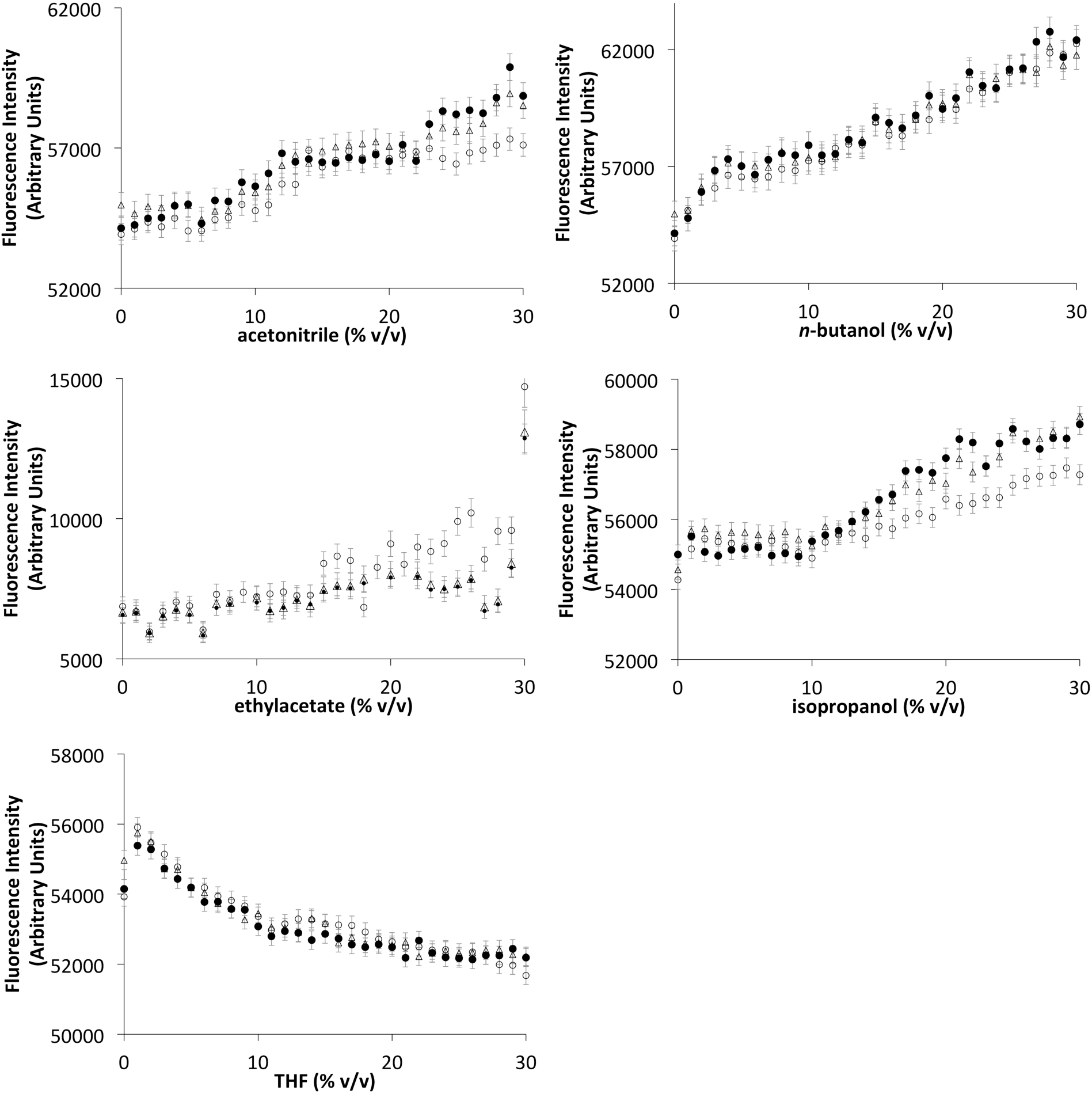
Fluorescence intensity measurements of apo-TK after incubation with increasing concentrations of organic solvents. Fluorescence intensity was measured at 340 nm with excitation at 280 nm. Apo-TK at 0.1 mg mL^−1^ in 25 mM Tris-HCl, pH 7.0 was incubated with (a) acetonitrile (b) *n*-butanol, (c) ethyl acetate (d) isopropanol (e) THF for (○) 1 hr, (Δ) 2 hr, (•) 3 hr at 25 °C prior to measurements. Error bars represent one standard deviation about the mean (n = 3).

**Fig. 9.**
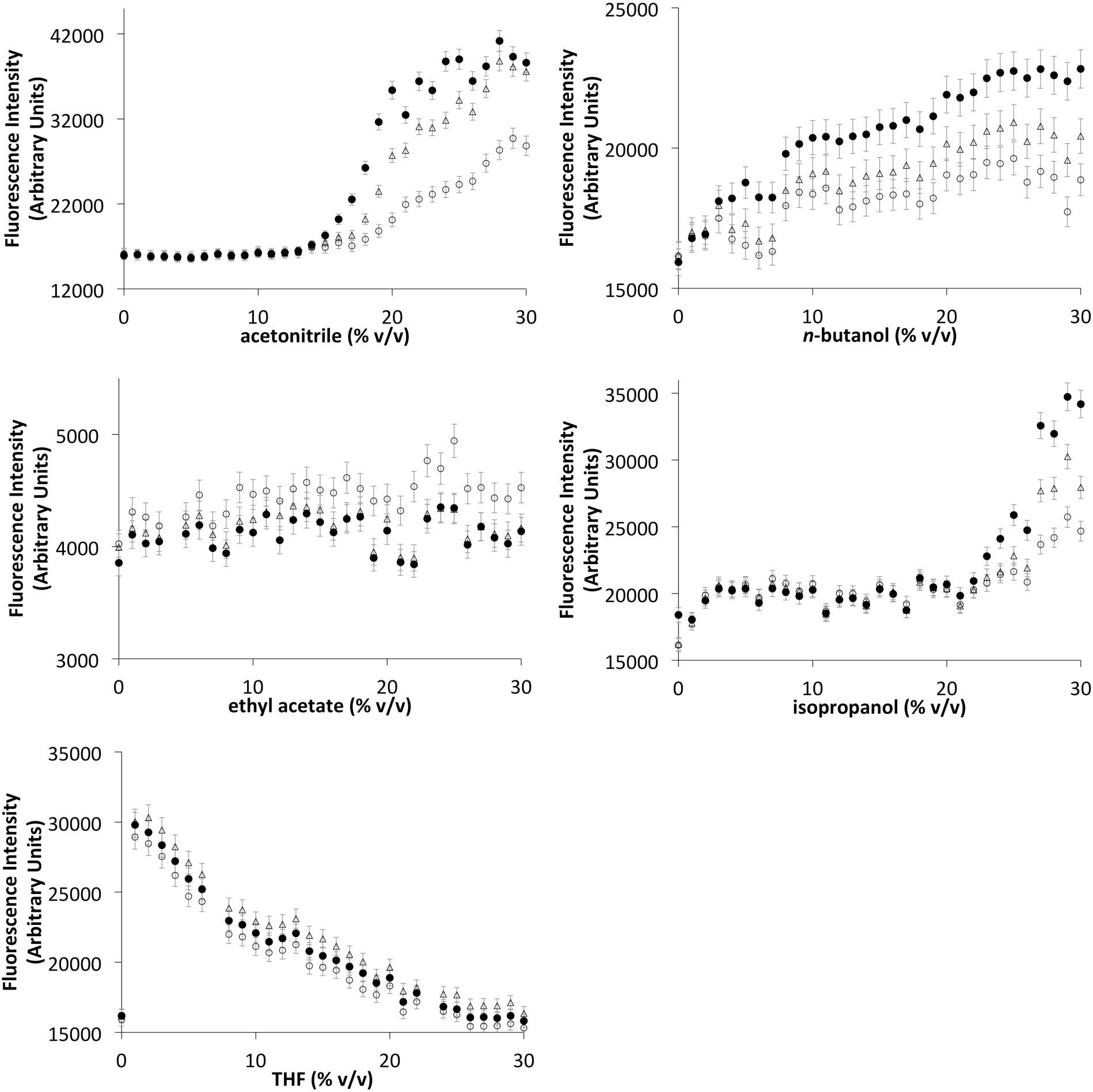
Fluorescence intensity measurements of holo-TK after incubation with increasing concentrations of organic solvents. Fluorescence intensity was measured at 340 nm with excitation at 280 nm. Holo-TK at 0.1 mg mL^−1^ in 5 mM MgCl_2_, 0.5 mM TPP, 25 mM Tris-HCl, pH 7.0, was incubated with (a) acetonitrile (b) *n*-butanol (c) ethyl acetate (d) Isopropanol (e) THF for (○) 1 hr, (Δ) 2 hr, (•) 3 hr at 25 °C prior to measurements. Error bars represent one standard deviation about the mean (n = 3).

For holo-TK the addition of co-solvents resulted in similar profiles to those for apo-TK, in EToAc and THF (Fig. 9). AcCN and iPrOH each gave no change initially and then increased sharply at above 15% (v/v) and 22% (v/v) respectively. In nBuOH, the fluorescence intensity showed a small increase that was monotonic overall, but with a small and sharp increase at 7-8%. These three sharp transitions are larger than any transitions observed in apo-TK, which didn’t aggregate, and they are consistent with the aggregation behaviour of holo-TK observed as time-dependent lag-phases in Fig. 7, and by DLS. Again, the lack of obvious transitions for EToAc and THF, and yet clear transitions at 15% (v/v) AcCN, 7-8% (v/v) nBuOH, and 22% (v/v) iPrOH, are consistent with the inactivation profiles for holo-TK in Fig. 3. However, the aggregate formation for holo-TK tended to occur at slightly higher solvent concentrations than the holo-TK inactivation, suggesting that it occurred *after* inactivation, and therefore that aggregates formed from an already inactive enzyme. This was consistent with the earlier observation by CD that inactivation was primarily due to a local unfolding event (or global unfolding in the case of AcCN with holo-TK), and that in some cases for holo-TK this also led to aggregation.

## 4. Conclusions

The deactivation of both apo-TK and holo-TK by co-solvents generally resulted from a local unfolding event that retained a near-native homodimeric structure. The only exception was for holo-TK in 20% (v/v) MeCN, which induced global unfolding. This is consistent with previous studies that suggest enzyme activity loss in polar solvents is primarily due to the stripping of water from the protein surface and enzyme active-site, which in turn affects the active-site geometry, stability and catalytic function (Prasad and Suguna, 2002; Simon et al., 2007). Local unfolding also led to aggregation for holo-TK in certain solvents, but only after enzyme deactivation had already occurred. For holo-TK in MeCN, enzyme inactivation was dominated by global unfolding.

The most hydrophobic co-solvents, with higher log(P) values, deactivated the enzyme at proportionally lower molar concentrations of co-solvent, although THF appeared to inactivate at lower concentrations than expected from this trend. The log(P) values also correlated well to the local unfolding rate observed by CD, indicating that the more hydrophobic solvents were better able to unfold the local structure responsible for enzyme deactivation.

Previous studies of other enzymes found that alcohols tend to disrupt tertiary structure and leave the secondary structure interactions largely undisturbed (Babu et al., 2001). The data for holo-TK were not consistent with this conclusion, as large changes in intrinsic fluorescence intensity, as observed for holo-TK in iPrOH or nBuOH, were correlated to aggregate formation, rather than significant loss of tertiary structure. More recently, the prolonged exposure to organic solvents was proposed to lead to a decrease in enzyme dynamics and active-site polarity, and also the reorientation of active site residues, which in turn affect the ionization state of the catalytic residues, and hence the stability of transition states and intermediates required for catalysis (Bansal et al., 2010). However, these mechanisms would be reversible equilibrium effects and so do not readily explain the loss of TK activity which was largely irreversible, as measured by activity assays after a 20-fold dilution out of the aqueous solvent mixture.

Interestingly, for THF, MeCN, and EtoAc, the more thermodynamically stable holo-TK did not retain more activity than the apo-TK after exposure to these co-solvents, indicating that solvent tolerance was not simply correlated to global conformational stability. However, the local unfolding rates were similar between apo-TK and holo-TK, and so it appears that the local unfolding measured by CD was not due to unfolding of the cofactor loops, as these are already unstructured in apo-TK. However, the structure of the cofactor loops, and/or presence of bound cofactors adversely influenced the retention of activity. Potentially, the unfolding of an adjacent local structure in both apo-TK and holo-TK, could occur with greater reversibility in apo-TK than in holo-TK. The 20-fold dilution into the reaction assay would then recover more activity for apo-TK than for holo-TK. Indeed, apo-TK unfolding was found previously to be more reversible than for holo-TK, when refolding TK from urea (Martinez-Torres et al., 2007).

It was also interesting that only MeCN induced global unfolding, and CD-observable aggregation, and then only in the holo-TK form. MeCN had one of the lowest log(P) values of the co-solvents tested, and so simple hydrophobicity of the solvent was not the reason. MeCN may be able to form additional interactions with the protein, not found for the other co-solvents, that enable it to unfold the protein more effectively. Alternatively, the apparent unfolding observed by CD may actually also have been due to aggregation, in which case the properties of MeCN promote more irreversible unfolding and aggregation than for the other solvents.

The ideal conditions for biocatalysis in industry require that the enzyme remains active for the duration of the biotransformation. Therefore, even the slow denaturation or aggregation by solvents, as observed for iPrOH, nBuOH and MeCN is potentially problematic for longer reactions, or for repeated enzyme use. Considering this, TK was most tolerant in 15% (v/v) (2.0 M) iPrOH, 10% (v/v) (1.9 M) MeCN, or 6% (v/v) (0.65 M) nBuOH as in these cases, the activity remained at >75%, whereas no aggregation was observed over 3 h.

Finally, this work has provided useful insights that will guide the future engineering of TK and other similarly complex enzymes. Improvements might be obtained by focussing enzyme engineering within the cofactor-binding loops and neighbouring regions close to the protein surface, as we have shown previously to be effective for the improvement of TK thermostability (Yu et al., 2017).

## Abbreviations

MeCN: Acetonitrile
CD: Circular dichroism
ERY: Erythrulose
DLS: Dynamic light scattering
EToAc: Ethyl acetate
GA: Glycolaldehyde
HPA: Hydroxypyruvate
iPrOH: Isopropanol
nBuOH: *n*-butanol
THF: Tetrahydrofuran
TFA: Trifluoroacetate
TK: Transketolase
TPP: Thiamine pyrophosphate
TPSA: Total polar surface area
[S_0_]: Concentration of solvent required for complete enzyme inactivation.

## Acknowledgements

The Royal Thai Government is acknowledged for the support of Phattaraporn Morris. The Mexican National Council for Science and Technology (CONACYT) is acknowledged for the support of Ribia García-Arrazola. The UK Engineering and Physical Sciences Research Council (EPSRC) is thanked for the support of the multidisciplinary Biocatalysis Integrated with Chemistry and Engineering (BiCE) programme (GR/S62505/01) at University College London (London, UK).

