## Supplementary Figure for "Biophysical characterization of the inactivation of *E. coli* transketolase by aqueous co-solvents"

### Appendix A. Supplementary data

***Fig. S1 (a-c).***

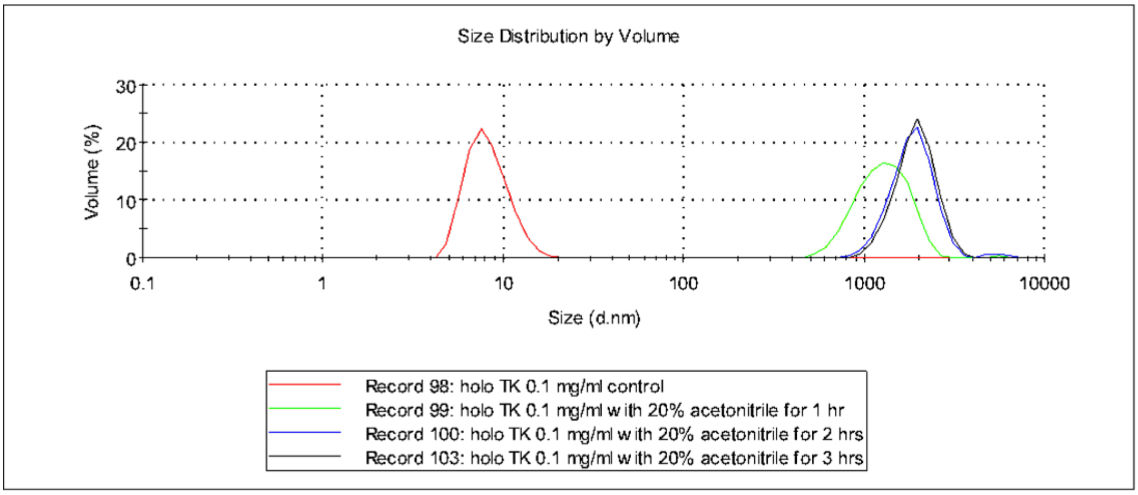


(a)

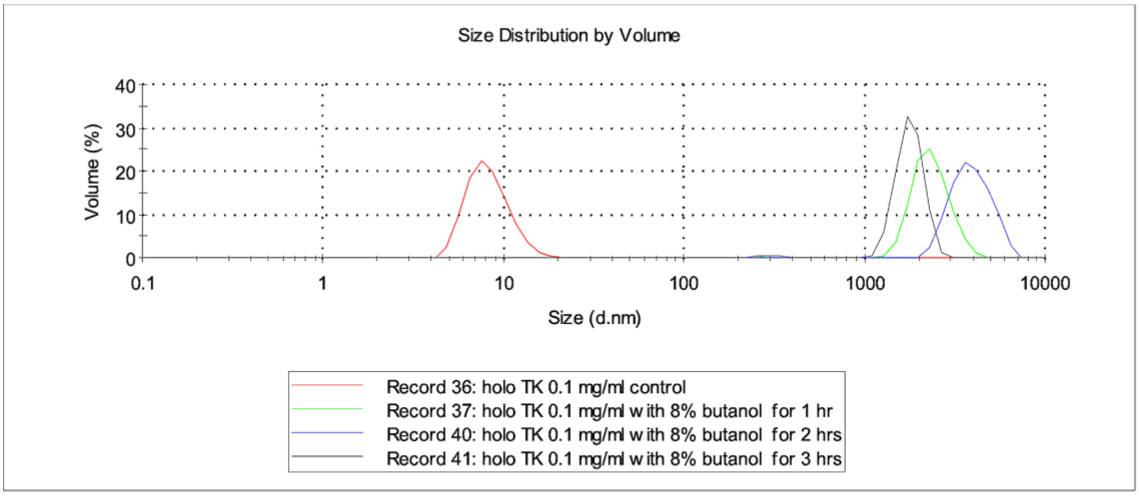


(b)

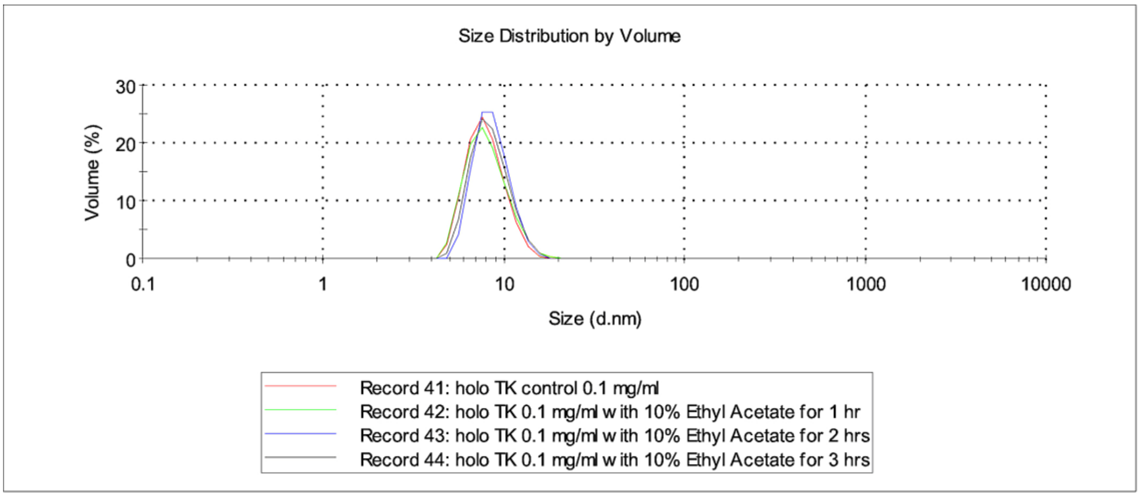


(c)

***Fig. S1 (d-e).***

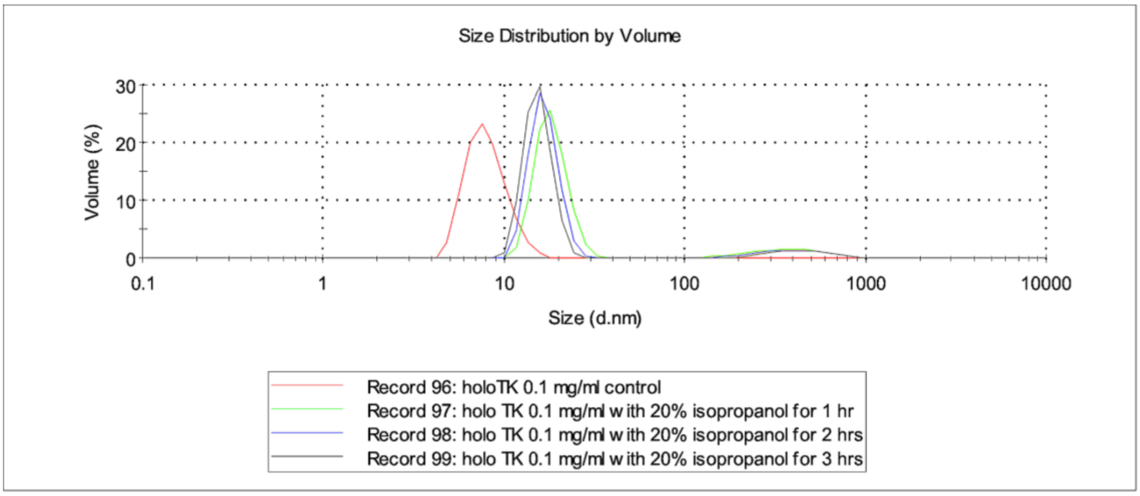


(d)

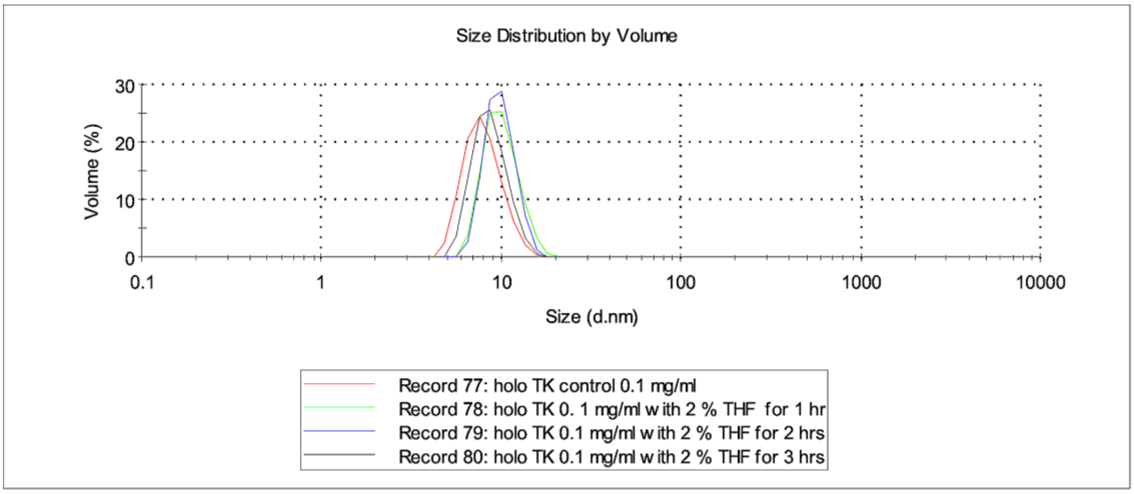


(e)

**Fig. S1.** Size distribution volume (%) of holo-TK estimated by dynamic light scattering (DLS) in the presence of organic co-solvents. Samples contained 0.1 mg $mL^{-1}$ protein in 25 mM Tris-HCl, pH 7.0, with cofactors (0.5 mM TPP and 5 mM $MgCl_2$), and (a) 20% acetonitrile, (b) 8% *n*-butanol, (c) 10% ethyl acetate, (d) 20% isopropanol or (e) 2% THF. Samples were incubated for 1, 2 and 3 h at 25 °C before measurement. Data from one replicate is shown for each sample type. Measurements were confirmed for triplicate samples.
